# The ventral hippocampal-nucleus accumbens shell circuit drives approach decisions under social novelty and learned cue approach-avoidance conflict

**DOI:** 10.1101/2024.06.27.600935

**Authors:** Dylan P Patterson, Nisma Khan, Kian Sassaninejad, Norman R Stewart, Emily Collins, Dylan Yeates, Andy CH Lee, Rutsuko Ito

## Abstract

Successful resolution of approach-avoidance conflict (AAC) is fundamentally important for survival, and its dysregulation is a hallmark of many neuropsychiatric disorders, and yet the underlying neural circuit mechanisms are not well elucidated. Converging human and animal research has implicated the anterior/ventral hippocampus (vHPC) as a key node in arbitrating AAC in a region-specific manner. In this study, we sought to target the vHPC CA1 projection pathway to the nucleus accumbens (NAc) to delineate its contribution to AAC decision-making, particularly in the arbitration of learned reward- and punishment signals, as well as innate signals. To this end, we used pathway-specific chemogenetics in male and female Long Evans rats to inhibit the NAc shell projecting vHPC CA1 neurons while rats underwent a test in which cues of positive and negative valence were presented concurrently to elicit AAC. Further behavioral assays of social preference and memory, reward and punishment cue processing, anxiety, and novelty processing were administered to further interrogate the conditions under which the vCA1-NAc shell pathway is recruited. Chemogenetic inhibition of the vCA1-NAc shell circuit resulted in animals exhibiting increased decision-making time and avoidance bias specifically in the face of motivational conflict, as the same behavioral phenotype was absent in separate conditioned cue preference and avoidance tests. vCA1-NAc shell inhibition also led to a reduction in seeking social interaction with a novel rat but did not alter anxiety-like behaviors. The vCA1-NAc shell circuit is therefore critically engaged in biasing decisions to approach in the face of social novelty and approach-avoidance conflict. Dysregulation of this circuit could lead to the precipitation of addictive behaviours in substance abuse, or potentiation of avoidance in situations of approach-avoidance conflict.

## Introduction

Approach-avoidance conflict resolution is a form of decision making that requires the effective evaluation of stimuli with mixed (positive and negative) valence. These opposing associations simultaneously incentivize an organism to approach and avoid the stimulus, creating a motivational conflict (1,2). The successful resolution of this conflict is not only necessary for survival but is also thought to be aberrant in mental health disorders. For instance, maladaptive approach biases are strongly evident in substance abuse patients and preclinical models of addictive behaviors, whereas avoidance biases are observed in individuals with anxiety and depression (3–7), underscoring the importance of elucidating the neural circuitry that supports approach-avoidance decision making.

A network of structures including the prefrontal cortex, striatum and limbic regions such as the hippocampus (HPC), amygdala, thalamus and hypothalamus has been implicated in mediating decision making under motivational conflict (8–14). In particular, the ventral aspect of the HPC (anterior in humans) has recently emerged from convergent human and animal research as a key node in the regulation of approach-avoidance conflict resolution, effecting approach and avoidance behaviors in a subfield-specific manner (15,16). Schumacher et al. (2018) found that GABAR-mediated inactivation of the vCA1 increased avoidance tendencies, while inactivation of the vCA3 increased approach tendencies in the face of motivational conflict, thus implicating the vCA1 in mediating cued approach behavior and the vCA3 in cued avoidance behavior(15). The divergent functions of these vHPC subregions in approach-avoidance conflict processing raises the question of whether this difference is mediated by their distinct extrinsic connections. To address this, we have recently identified the extrinsic connectivity of the vCA3 with the lateral septum, the sole target of long range vCA3 projections, to be critical in mediating cued avoidance behavior under conflict(17).

The vCA1, unlike the vCA3, has extensive connectivity downstream (18), among which, the strong connectivity between the vCA1 and nucleus accumbens (NAc) is of particular interest in modulating approach-avoidance behaviors in the face of motivational conflict (19). Traditionally, the NAc has been thought to mediate reward-directed behavior and act as a limbic-motor interface that functionally links motivation and action(20). However, more recent evidence suggests the NAc is involved in the evaluation and integration of both positively and negatively valenced information and their influence over behavioral output in a regionally specific manner (21–26). For instance, NAc shell GABA microcircuits are thought to represent appetitive to aversive information along a rostro-caudal gradient (17–18). The NAc core is implicated in mediating cued influences over pavlovian or goal-directed behavior (27), and the caudal core GABA has also been implicated in promoting an avoidance bias during exposure to a cue inducing approach-avoidance conflict (14).

Glutamatergic projections from the vCA1 to the NAc predominantly target the shell, and this pathway has been implicated in promoting stress-induced negative affective/motivational states (28), inhibiting feeding (29), as well as promoting positively motivated behaviors such as social (30) and contextual (31) reward-seeking. However, the role of NAc shell-projecting vCA1 in guiding approach-avoidance choices when faced with stimuli of opposing valence concurrently remains unexplored. To address this, animals underwent chemogenetic inhibition of the vCA1-shell pathway while performing a cued conflict test in which they were given a choice to explore a learned cue imbued with both appetitive and aversive qualities (to induce conflict), and a cue associated with no outcome. We report that inhibiting vCA1 glutamatergic afferents to the NAc shell disrupts evidence accumulation in the face of a conflict decision, revealing a hitherto undiscovered role of a vCA1–NAc shell in approach-avoidance conflict resolution.

## Results

Histological verification of viral expression and injector placement revealed viral transduction within the vCA1 (and sometimes vCA3) regions of the vHPC for both hM4Di-mCherry and GFP constructs. Terminal expression of the constructs was found to be confined to the NAc shell, with cannula and infuser tip placement verified to be located within the NAc shell to enable targeted inhibition of NAc-projecting vCA1 neurons (Figure 1A-C). Assessment of cfos positive (+) cell density in the NAc shell revealed that microinfusion of CNO induced a significant decrease in cfos+ cells in the shell of hM4Di-mCherry expressing animals, compared to those microinfused with saline (Figure 1D, F*drug* (1,25) = 11.91, p= .002, η^2^ =.32, F*sex* (1,25) = 3.28, p = .082, F*drug* x *sex* (1,25) = 1.48, p = .24), thereby verifying the dampening of activity of NAc shell-projecting vCA1 neurons.

**Figure 1.**
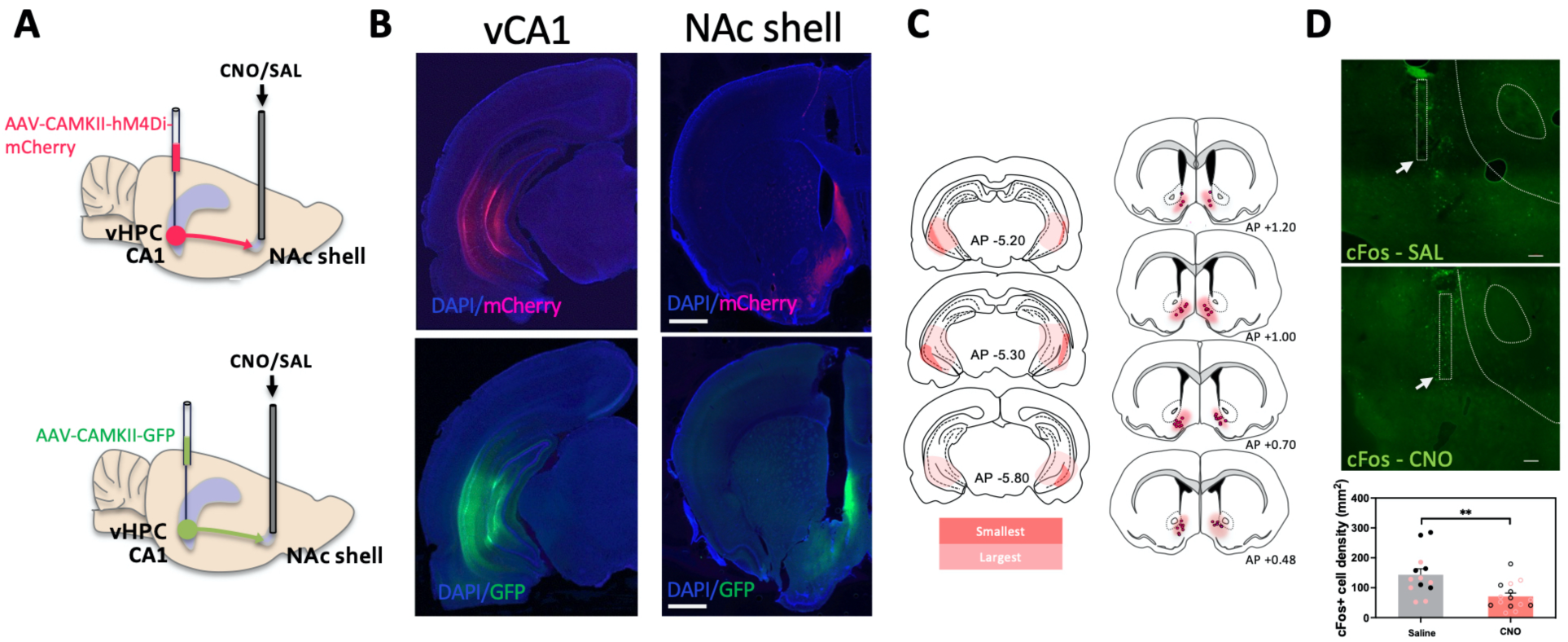
Chemogenetic inhibition of ventral hippocampus CA1 (vCA1) to nucleus accumbens (NAc) shell projections. **A** Schematic diagram of inhibitory DREADDS (hM4Di-mCherry) or empty vector (GFP) transduction in vCA1 and cannula implantation in NAc shell via which clozapine N oxide (CNO) or saline was microinfused for pathway-specific inhibition. **B** Representative images of somatic transduction of hM4Di-mCherry or empty vector GFP in vCA1 (left) and axonal transduction in the NAc shell (right). Scale bar, 1mm **C** Cannula placements in the NAc shell (DREADDs group only) and the minimum and maximum DREADDs expression in vCA1 and NAc shell, AP: Anterior-Posterior to bregma. **D.** Representative images of cFos expression in the NAc shell following either saline (SAL) or CNO (1mM) infusion, showing cannula tract, and tip (arrow). Scale bar, 100um. Intracerebral CNO infusions attenuated cFos expression in areas of the NAc shell that received hM4Di positive inputs (bottom graph). Red symbols depict female data. Data show means ± SEM. ** p < .01.

Animals were first trained to associate three visual-tactile cues with distinct outcomes: appetitive, neutral and aversive in the radial maze (Figure 2A). Every four conditioning sessions, learning was assessed using a conditioned cue acquisition test in which animals were allowed to freely explore the cues for 5 minutes in the absence of any reinforcement (Figure 2B). All animals successfully acquired cue valence by their final (second or third) acquisition test (Figure 2B, F*cue*(1.71, 49.15) = 160.99; p < .001) and spent significantly more time exploring the appetitive arm compared to the neutral arm (p < .001) and significantly less time exploring the aversive arm compared to the neutral arm (p < .001). There were no significant differences in the cue acquisition pattern between sex (male vs. female) virus (GFP vs hM4Di) and future drug (saline vs CNO) groups (F*sex* x *virus x drug x cue* (1.705,86.93) = 0.14; p = .87).

**Figure 2.**
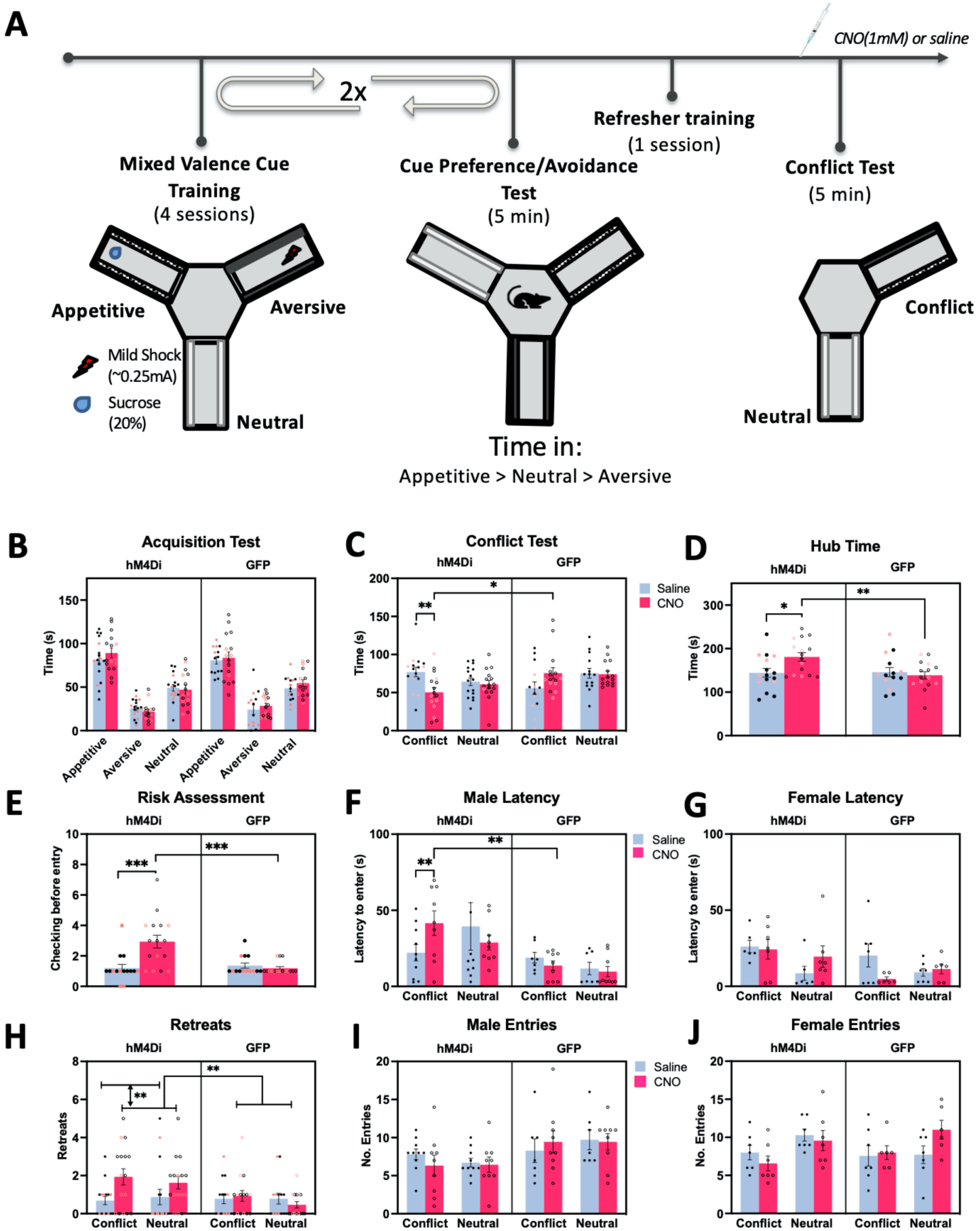
Effect of chemogenetic inhibition of vCA1-NAc shell on approach-avoidance conflict. **A**. Experimental timeline showing mixed valence cue training (2 x 4 sessions), followed by a cue preference/avoidance test (5min), refresher training session, and finally the cue conflict test, prior to which animals received intracranial microinfusion of CNO (1mM) or saline. **B**. After two rounds of 4 conditioning sessions, animals (hM4Di Sal: n=17, hM4Di CNO: n=16, GFP Sal: n=14, GFP CNO: n=16) successfully acquired cue-outcome associations, demonstrating a greater amount of time in the appetitive vs the neutral arm (p < 0.001) and less time in the aversive vs the neutral arm (p < 0.001). **C.** In the conflict test, rats were given a choice to explore a neutral cued arm or an arm with the appetitive and aversive cues superimposed (conflict cue). Time spent in the conflict cued arm was significantly reduced in the vCA1-NAc shell inhibited group (hM4Di CNO) compared to control groups (hM4Di SAL, GFP). Red symbols depict females. **D.** Amount of time spent in the hub (central compartment), and **E.** Risk assessment (checking) behavior were also significantly increased in the vCA1-NAc shell inhibited group, compared to control groups. **F-G.** Latency to enter the conflict cue arms for the first time was increased in the males but not females after vCA1-NAc shell inhibition. **H**. Number of retreats in both conflict and neutral arms were more pronounced in the vCA1-NAc shell inhibited group compared to other control groups. **I-J** vCA1-NAc shell inhibition did not alter arm entries, however, the females made more entries into the neutral than conflict arms. Data show Means ± SEM. * p < .01, ** p <.01, ***p <.001

Following successful cue acquisition, animals underwent a conflict test (Figure 2C), in which they were given a choice to explore an arm with the neutral cue or an arm with both the appetitive and aversive cues superimposed (conflict arm) under extinction conditions. Chemogenetic inhibition of the vCA1-NAc shell shell pathway significantly and selectively reduced the time spent exploring the conflict cue compared to the hM4Di saline (*p* = .007) and GFP CNO controls (*p* = .017, Figure 2C, F*virus x drug x cue* (1, 53) = 4.81; p = .033, η^2^ = .083), irrespective of sex (F*virus x drug x cue x sex* (1, 53)= 1.45, p= .23). However, all animals spent equal amounts of time exploring both the conflict and neutral arms (all p > .10), indicating that animals did not have an approach or avoidance bias for the conflict cue over the neutral cue. Nevertheless, a separate examination of the central hub time revealed that chemogenetic vCA1-NAc shell inhibition caused a significant increase in the time spent within the hub compared to saline (p= .019) and GFP CNO control rats (p = 0.005, F*virus x drug* (1, 53) = 4.06; p = .049, η^2^ = .071, Figure 2D). No significant differences between the hub time of GFP saline and CNO animals were observed (p = 0.65).

To further examine the increased hub time coupled with the decreased conflict arm time in the CNO-infused hM4Di group, we also analyzed risk-assessment behavior, which was defined as the rats looking back and forth between the two arms before committing to their first entry into an arm. CNO-infused hM4Di animals exhibited significantly more ‘looking’ behavior than saline (p = .001) and GFP CNO controls (p < .001, Figure 2E, F*virus x drug* (1, 53) = 7.47; p = .008, η^2^ = .12) (Figure 2E). The latencies to enter into the conflict and neutral arms for the first time were also analyzed, and we found that CNO-infused hM4Di males, but not females, were significantly slower to make entries into the *conflict* arm compared to saline-infused hM4Di male (p = .012) and GFP CNO male (p < .001) controls, as well as their female CNO-infused hM4Di counterparts (p < .001, *Fsex x virus x drug x cue* (1, 53) = 4.28; p = .043, η^2^ = .075 Figure 2F-G).

In addition, we analyzed the number of retreats exhibited by the rats once they entered the arms as an index of aversion. CNO-infused hM4Di animals exhibited significantly more retreats overall (irrespective of cue type or sex), compared to saline-infused hM4Di (p = .002) and GFP CNO (p < 0.004) controls (F*virus x drug* (1, 53) = 5.87; p = .019, η^2^ = .10, F*virus x drug x cue x sez* (1, 53) = 1.85; p = .18 Figure 2H). Finally, we also analyzed the total number of arm entries during the conflict test and found significant sex differences, in that female rats entered the neutral-cued arm more frequently than the conflict-cued arm (Figure 2I-J, F*sex x cue*(1, 53) =6.83; p = .012, η^2^ = .11), albeit CNO induced inhibition of the vCA1-NAc circuit did not induce any significant changes in the number of entries into the conflict and neutral arms (all interactions involving drug, p > .13).

Next, to assess if vCA1-NAc shell inhibition influenced cued approach-avoidance decision in the *absence of motivational conflict*, conditioned cue preference (CCP) and conditioned cue avoidance (CCA) tests were conducted separately (Figure 3A, B & D). All animals spent significantly more time in the appetitive arm than the neutral arm in the appetitive cue preference test (Figure 3C, F*cue* (1, 51) = 63.38; p < .001, η^2^ = .55), irrespective of virus, drug group or sex (*Fsex x virus x drug x cue* (1, 51) = .18; p > .68). All animals also spent significantly less time in the aversive arm than the neutral arm in the aversive cue test (F*cue* (1, 51) = 107.87; p < .001, η^2^ = .68). There was evidence of CNO-infused hM4Di males spending less time in the neutral arm than the CNO-infused GFP male controls (p = .045), but not when compared to saline-infused hM4Di (p = .19) group (post-hoc tests conducted after significant interaction; *Fsex x virus x drug x cue* (1, 51) = 4.06; p =.049, η^2^ = .074, Figure 3E). The time spent in the hub during CPA was significantly higher in the CNO-infused hM4Di rats compared to CNO-infused GFP rats (p = .002, F*virus x drug* (1,51)= 4.03, p = .05, Figure 3F), but not saline-infused hM4Di rats (p =.096). However, hub time during the CPP was not significantly different between the CNO or saline-infused hM4Di or GFP rats (all tests with Drug, p > .34, Figure 3G). Latencies to enter the arms during CPP and CPA were not significantly different following CNO infusions in hM4Di-expressing rats (all tests involving Drug, p > .29 for CPP, p > .16 for CPA, Figure 3H-I), with all rats taking longer to enter the aversively cued arm (F*cue* (1, 51) = 9.75; p = .003, η^2^ = .16), but no significant difference in latencies to enter the appetitively vs. neutral cued arms was found (F*cue* (1,51) = 1.59; p = .21 η^2^ = .03). As expected, the number of retreats was significantly higher in the aversively cued arm vs. neutral arm in all animals irrespective of virus, drug or sex (F*cue* (1, 51) = 37.31; p < .001, η^2^ = .42, all other tests p > .08, Figure 3J) and retreat behavior was never observed during CPP (all 0 - data not shown). Risk assessment behavior was observed significantly more during CPA than CPP (F*test* (1,51)=7.59, p =.008, η^2^ = .13, Figure 3K,L), with no significant difference in the performance between CNO and saline-infused hM4Di or GFP rats (all tests with drug, p > .30). Finally, rats made more entries into the appetitively cued arm compared to neutral arm (F*cue* (1, 51) = 10.25; p = .002, η^2^ = .17, Figure 3M) and less entries into the aversively cued arm (F*cue* (1, 51) = 97.3; p < .001, η^2^ = .66, Figure 3N). Altogether, these results indicate that chemogenetic inhibition of the vCA1-NAc shell pathway did not affect conditioned cue preference or avoidance behavior *per se*, and that the observed effects of vCA1-NAc shell inhibition were specific to the conflict test.

**Figure 3.**
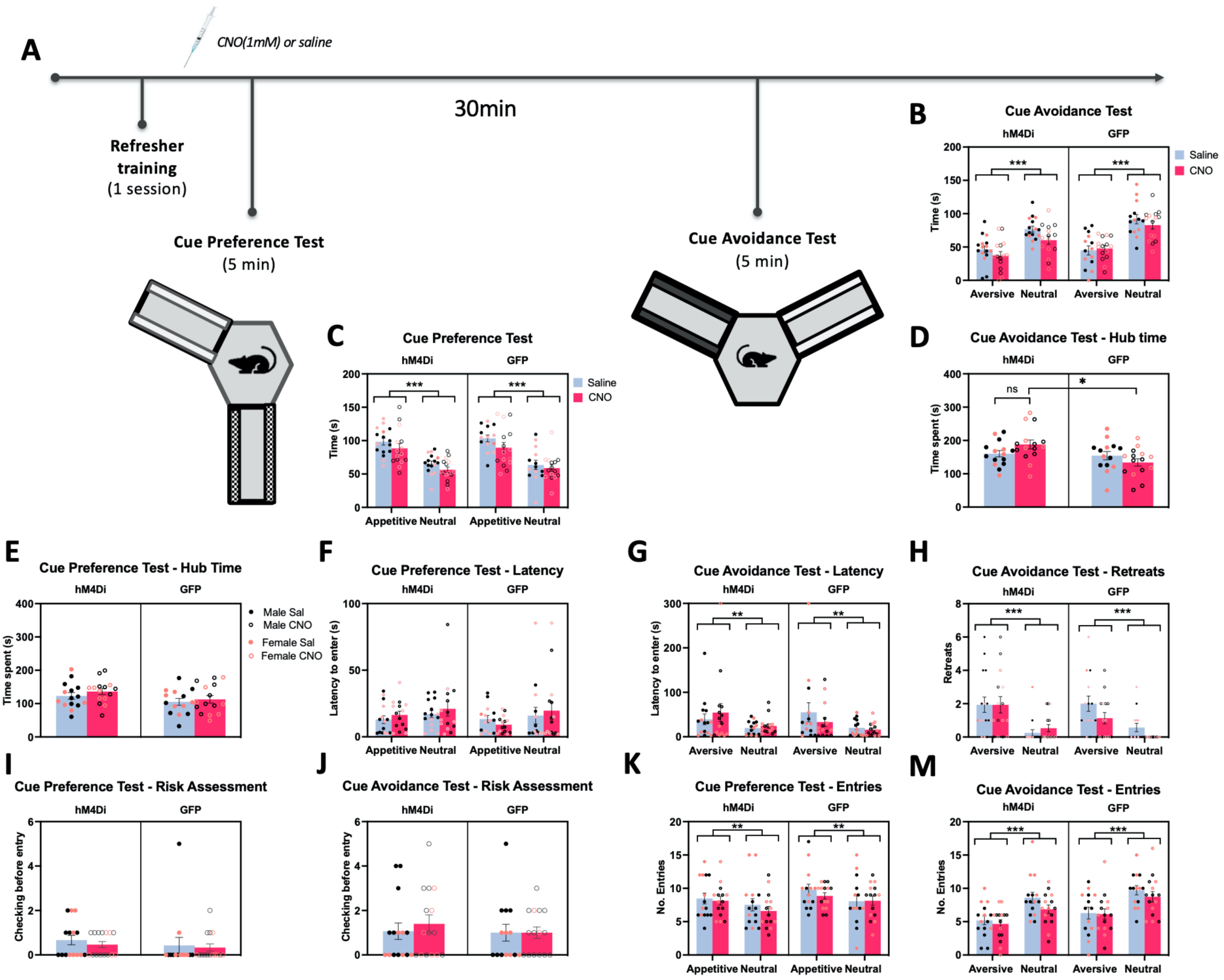
No effect of vCA1-NAc shell inhibition on cue preference and avoidance tests. **A.** Experimental timeline showing the administration of a refresher conditioning session following the conflict test, then the administration of cue preference (appetitive vs. neutral cues) and cue avoidance (aversive vs. neutral cues) tests in counterbalanced order, prior to which animals were microinfused with CNO or saline. **B,C.** All animals (hM4Di Sal: n=16, hM4Di CNO: n=15, GFP Sal: n=14, GFP CNO: n=15, note attrition of two rats after the conflict test due to blocked cannnula) spent significantly less time in the aversive vs. neutral arm in the cue avoidance test, and spent significantly longer in the appetitive vs. neutral arm in the cue preference test, irrespective of pathway inhibition. **D,E.** Time spent in the hub (central compartment) during the cue avoidance test was significantly greater in the vCA1-NAc shell inhibited group compared to the GFP CNO-infused group but not hM4Di SAL infused group. There was no difference in the hub time during the cue preference test between any groups**. F,G.** Latency to enter the aversive cued arm for the first time was significantly higher compared to latency to enter the neutral arm in all groups. No difference in latency to enter the appetitive vs. neutral arms was observed. **H.** The number of retreats observed in the aversively cued arm was significantly higher than in the neutral cued arms in the cue avoidance test. **I,J** There was no difference in risk assessment behavior between all groups in cue preference and avoidance tests. **K,M.** All animals spent made more entries into the appetitive arm over the neutral arm in the cue preference test, and made less entries into the aversive arm over the neutral arm in the cue avoidance test, irrespective of pathway inhibition. Data show Means ± SEM. * p < .01, ** p <.01, ***p <.001

We also conducted a cue novelty preference test in a subset of animals, in which a novel set of cues was presented in one arm of the Y-maze, and the neutral cues in another arm to assess whether the observed effects in the conflict test were a function of the relative novelty of the conflict cue. Novelty cue preference was evident in both CNO injected hM4Di and GFP groups, with animals spending more time exploring the novel cues over the neutral cues over the course of 3min, then 5min (F*arm* (1,10) = 8.93; p = .02, η^2^ = .43, no significant effects of *virus*, all p > .43, Figure 4B). No significant differences were observed between the hM4Di and GFP groups in hub time (F*virus* (1,10) = 1.18; p = .30, not shown), total number of retreats (F*virus*(1,10) = 3.00, p = .11, not shown), risk assessment ‘looks’ (F*virus*(1,10) = 3.64, p = .09, not shown), nor in the latency to enter either of the arms (F*virus*(1,10) = 1.30, p = .28).

**Figure 4.**
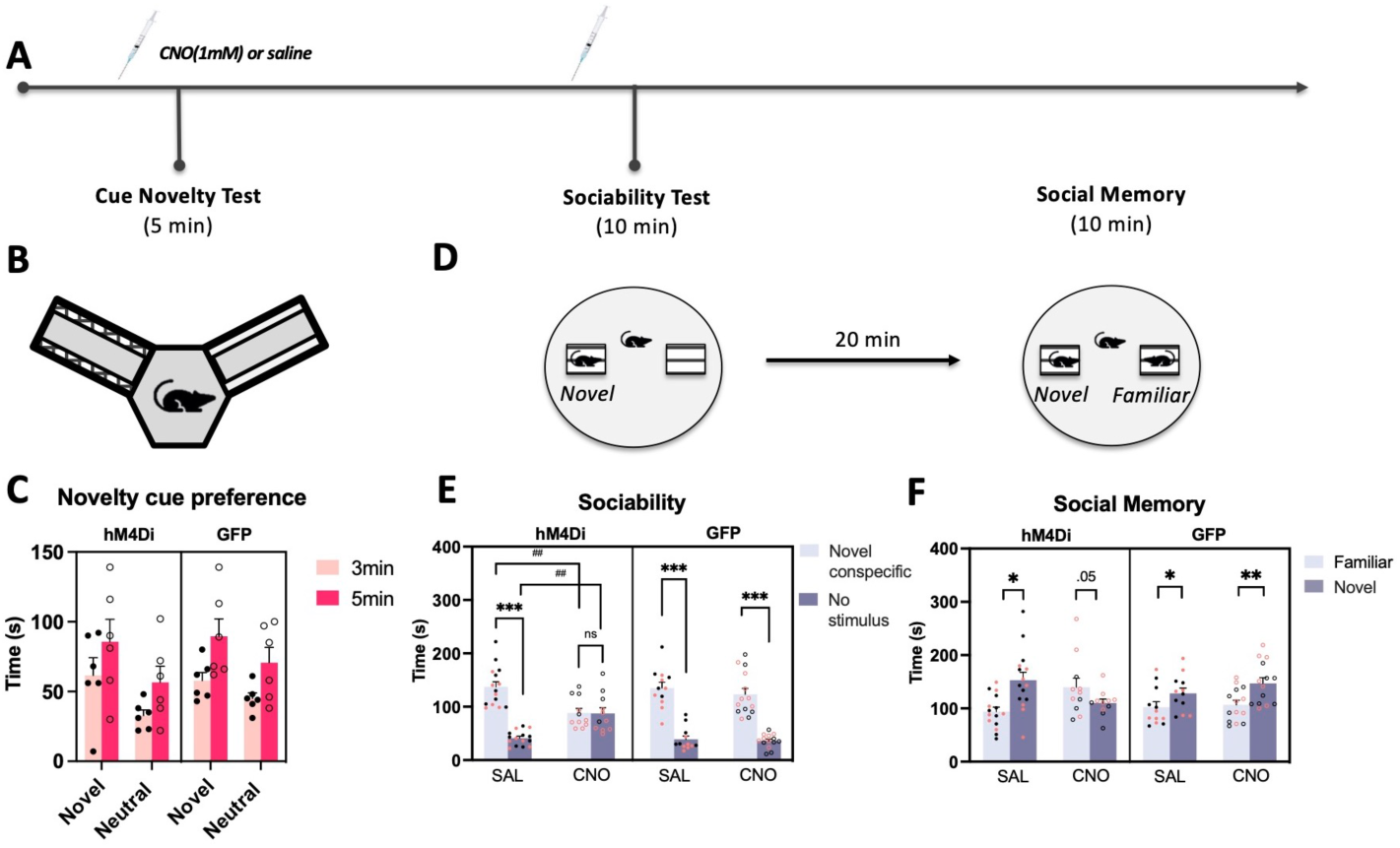
Effect of vCA1-NAc shell inhibition on assays of novelty processing, social seeking and discrimination. **A.** Timeline of experimental procedures; following the completion of the cue preference/avoidance tests, animals underwent testing in: cue novelty test, social interaction and novelty seeking tests. Prior to each of the tests, animals were microinfused with CNO or Saline**. B.** Animals (hM4DI CNO: n=6, GFP CNO: n=6) spent more time engaging with novel over neutral cues in the Y maze, regardless of construct group (hM4Di & GFP rats all microinfused with CNO). **E.** vCA1-NAc shell inhibited animals spent significantly less time interacting with the novel conspecific over an empty cage compared to control groups, and failed to show preference for social interaction (hM4Di Sal: n=15, hM4Di CNO: n=11, GFP Sal: n=12, GFP CNO: n=14, note n=8 were excluded due to data collection error). **F**. vCA1-NAc shell inhibited animals spent more time interacting with the familiar rat and significantly less time interacting with (i.e., avoidance of) the novel rat compared to controls. Data show Means ± SEM. * p < .01, ** p <.01, ***p <.001, ^##^ p < .01

Since the vCA1-NAc shell circuit has also been implicated in the retrieval of social memory, a two phase social discrimination task was conducted whereby animals were first given the opportunity to interact with a novel caged conspecific versus an empty cage (sociability test), and then subsequently given the choice to interact with the familiar caged conspecific from Phase 1, or a novel caged rat (social memory test, Figure 4D). Chemogenetic inhibition of the vCA1-NAc shell pathway significantly altered social-seeking behavior in the sociability test (Figure 4E, F*virus x drug x stimulus* (1, 44) = 19.45; p < .001, η^2^ = .31), with CNO-infused hM4Di animals spending significantly less time interacting with the novel caged conspecific compared to saline-infused hM4Di (p = .004) and CNO-infused GFP control animals (p = .045), and significantly increased time interacting with an empty cage compared to controls (hM4Di Sal: p <.001, GFP CNO: p < .001). Furthermore, whereas all three control groups exhibited significantly higher exploration of the caged conspecific compared to the empty cage (no stimulus, all p<.001), vCA1-NAc shell inhibited animals failed to show this discrimination (p= .97).

vCA1-NAc shell pathway inhibition significantly altered social memory test performance (Figure 4F, F*virus x drug x novelty* (1, 43) = 12.98; p < .001, η^2^ = .23). CNO-infused hM4Di animals spent significantly more time with the familiar rat (p = .01) and significantly less time with the novel rat (p = .003) compared to saline controls. Moreover, unlike the other control groups, CNO-infused hM4Di animals failed to exhibit preference for the novel over familiar rat, and instead showed marginal avoidance of the novel rat (hM4Di CNO: p = .053, hM4Di Sal: p <.001, GFP CNO: p = .002, GFP Sal: p=.006). The drug (CNO) had no effect on overall interaction time during the test (F*virus x drug*(1, 44) = 0.37; p = .37, F*drug*(1, 28) = 0.36; p = .55).

To confirm that the decreased conflict cue exploration and increased latency to enter during the conflict test were not due to a general decrease in movement, a locomotor task was conducted. Overall, there were no significant effects of CNO on locomotor activity (all drug effects, p > .054, Figure 5B,C), with habituation (reduction) of locomotor activity over the 60 min session, as expected (F*bin* (6.12,317.99) = 45.57; p < .001, η^2^ = .47). There were significant sex differences, however, with the females displaying elevated locomotor activity across the time bins compared to males (F*sex* (1,52)= 5.60, p= .022, η^2^ = .097, F*binxsex* F(6.12,317.99) = 2.86, p = .01, η^2^ = .05).

**Figure 5.**
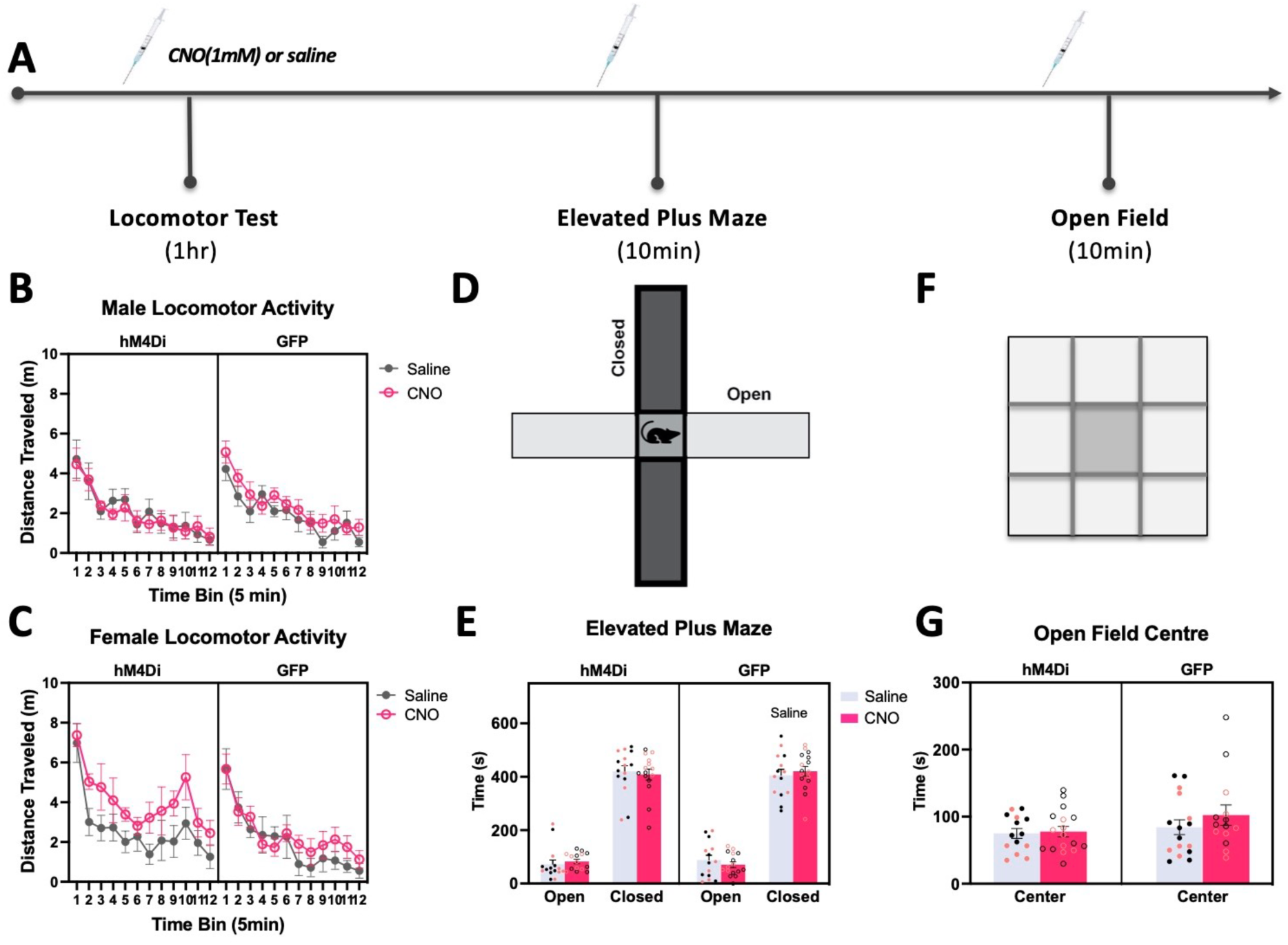
Effect of vCA1-NAc shell inhibition on assays of locomotor activity and innate anxiety. **A.** Timeline of final set of experimental procedures involving locomotor activity test, elevated plus maze, and finally, open field test. Prior to each of the tests, animals were microinfused with CNO or Saline **B,C.** Females showed elevated locomotor activity overall compared to males, but this was irrespective of vCA1-NAc shell inhibition. **D,E.** All animals spent significantly more time exploring the closed vs. open arms of the elevated plus maze with no group differences. **F,G.** All groups spent significantly more time exploring the periphery of the open field vs the middle (p < .001). Data show Means ± SEM.

Additionally, to rule out alterations in innate anxiety as an explanation for the increased latency and decreased conflict exploration, both elevated plus maze and open field tests were conducted. Animals spent significantly more time exploring the closed arms of the EPM (p < .001, Figure5D&E, F*arm*(1, 50) = 376.06; p < .001, η^2^ = .88) and the open field periphery (p < .001, Figure 4F&G, F*area*(1, 25) = 879.162; p < .001). However, there were no significant differences in elevated plus maze (F*virus x drug x arm*(1, 50) = .5 ; p = .48) nor open field test (F*virus x drug x area*(1, 25) = .704; p > .05) behavior between virus or drug groups. A separate examination of EPM middle time revealed that chemogenetic vCA1-NAc shell inhibition had no effect on time spent within the middle area compared to saline controls (t(13) = 0.655; p > .05, data not shown). Additionally, no significant differences in middle time were seen between GFP control animals (t (14) = 0.655; p > .05).

Together, our results indicate that vCA1-NAc shell inhibition resulted in decreased conflict cue exploration compared to controls. This behavior was accompanied with increased hub time, retreat behavior, latency to enter (in males) and risk assessment behavior, which is together indicative of potentiated avoidance bias in the face of approach-avoidance conflict. Furthermore, these results were specific to the conflict test as they occurred independently of alterations in locomotion, anxiety or cue avoidance/preference/novelty. Our results also reveal that vCA1-NAc shell inhibition impaired sociability and social novelty preference.

## Discussion

In the present study, chemogenetic inhibition of the vCA1-NAc pathway resulted in a decision-making deficit and a potentiation of avoidance bias in the face of motivational conflict, as evidenced by an increased time spent in the hub, enhanced latency to enter in the conflict cued arm (in males only), and potentiated retreat and risk assessment behavior prior to entering the conflict cued or neutral cued arm. We further demonstrated that these effects occur specifically under conditions of cue-elicited conflict, as vCA1-NAc inhibition did not impact any indices of decision making or approach/avoidance behaviors in the absence of conflict (in conditioned cue preference/avoidance tests). Moreover, the observed effects were independent of any alterations in novelty processing, locomotion, or anxiety. Interestingly, in seeking to replicate previous findings of impaired social discrimination elicited by vCA1-NAc inhibition, we found that vCA1-NAc-inhibited animals were fundamentally impaired in social interaction- and novelty-seeking. Together, our results implicate the vCA1-NAc shell in driving the decision to approach in scenarios that involve motivational conflict or social novelty, revealing a nuanced role for the circuit in supporting approach avoidance decisions under specific conditions.

We hypothesized that inhibition of the vCA1-NAc shell pathway would result in increased avoidance behavior in the face of an approach-avoidance conflict, in line with previous findings showing that pharmacological inhibition of NAc D1R neurons, the caudal NAc core and the vCA1 resulted in an increased avoidance of the conflict cue, characterized by a significant reduction in time exploring the conflict arm vs the neutral arm in the same task(14,27,33). Consistent with this hypothesis, we observed that vCA1-NAc shell-inhibited animals exhibited a conflict avoidant phenotype overall, and additional traits indicative of indecision. These animals spent significantly less time exploring the conflict arm compared to all other control groups. They retreated more frequently once in the conflict cued arm, indicating a heightened state of aversion in the conflict test. Further behavioral analysis revealed that vCA1-NAc shell inhibited animals exhibited significantly increased central hub time, increased time deliberating before making a decision to enter an arm, and were more latent and hesitant to make an initial entry into an arm, altogether pointing to a decision-making deficit. Moreover, vCA1-NAc shell inhibition failed to induce the same behavioral pattern during conditioned cue preference and avoidance tests, as well as novelty cue preference test, suggesting that the decision-making deficit was specific to the presence of a conflict scenario. Thus, the role of NAc shell-projecting vCA1 neurons in cued approach-avoidance conflict processing is more complex than originally hypothesized, and our results point to a potential role of this pathway in the facilitation of approach during decision making under motivational conflict. Indeed, this role is consistent with previous reports of elevated NAc shell dopamine being linked to rats that were classified as risk-taking, consistently choosing a larger reward paired with high probability of concomitant shock delivery over a certain, small reward (34) as well as NAc shell inhibition inducing risk-aversion (reduction in choice of larger reward with lower probability of delivery) in a probabilistic risk discounting task (35). Interestingly, the human NAc shell exhibits increased activity only when participants choose to accept (aka approach) an offer with a 50/50 chance of gaining or losing money in a monetary gambling task (36). Altogether, vCA1 afferents to NAc shell may be important in biasing decisions towards ‘approach’ in times of reward uncertainty and conflict.

In accord with previous findings, we observed impaired social discrimination in animals with vCA1-NAc shell circuit inhibition, evidenced by the failure to exhibit a preference for a novel, over familiar rat. Previous work has shown that manipulations of the vCA1, and its projections to the NAc induce similar social memory impairments (37,38). However, in light of the current findings that vCA1-NAc inhibited rats exhibited a lack of preference for social interaction over a non-social stimulus in the sociability test, an alternative interpretation to the prevailing mnemonic explanation for these social discrimination impairments may be that vCA1-NAc inhibition diminishes the drive to seek the novel conspecific, which may be perceived as a conflict stimulus with inherent rewarding and aversive properties. Indeed, in the present study, we observed an effect bordering on avoidance of the novel rat, similar to a previous finding in which an avoidance of a novel conspecific was reported following DA receptor antagonism of the NAc shell combined with cannabinoid agonism of the vHPC (39). Interestingly, vCA1-NAc inhibition did not impact the detection of a novel set of bar cue stimulus in the Y-maze in the present study, suggesting that the presence of a motivational conflict involving the evaluation of the potential of reward versus punishment may be a prerequisite of vCA1-shell engagement. Relatedly, despite the abundance of data showing that vHPC lesions induce anxiolytic effects (32,40–42), we did not observe alterations in the performance of EPM or open field tests following vCA1-shell inhibition. These ethological tests of anxiety represent different forms of approach-avoidance conflict scenarios wherein animals must decide between approaching safe, enclosed spaces or venturing into open areas that could potentially be dangerous. Of further note is that our findings are consistent with previous reports showing that vCA1-NAc shell circuit manipulation fail to perturb anxiety-related behavior, as measured by ethological tests (28,30).

The vHPC-shell pathway has been previously implicated in promoting stress-induced negative affective (depressive-like) states such as anhedonia and learned helplessness (28), susceptibility to chronic social stress, and inhibition of ethanol-seeking(43), suggesting that stress- or alcohol-induced maladaptive behavior may be mediated by an aberrant engagement of this pathway (but see ^26^ for evidence of inhibition of food seeking in stress-naïve animals). However, the vHPC-shell pathway has also been shown to support positively motivated behavior. For instance, LTP induction in the pathway promotes conditioned place preference in stress- and drug-naïve animals (31). Similarly, optogenetic stimulation of the vHPC-shell circuit induces conditioned place preference, and supports instrumental responding (44). Furthermore, as discussed above, vHPC-shell activation promotes pro-social behavior (32–34). Thus, the vHPC-shell pathway can mediate both negatively and positively motivated behavior under different circumstances, and the present findings extend this body of work by demonstrating that under situations in which stimuli of contrasting valences are presented simultaneously, the vCA1-shell pathway promotes approach. The mechanism underlying this seemingly dichotomous function of the vHPC-shell circuit in promoting approach and avoidance behaviors under varying conditions is unclear. However, one possibility is that the nature of behavioral output (approach or avoid) is determined by the activation pattern of the NAc along the rostral-caudal axis, which is known to be functionally distinct, with the rostral aspect regulating appetitively motivated behaviors and caudal aspect aversively motivated behaviors (17–22,24). Manipulations that reduce afferent input activity in the rostral NAc typically increase reward-seeking and consumption(29) , whereas increase in activity in this region has the opposite effect of reducing food seeking. Calcium imaging evidence also points to divergent activity in rostral (decrease) vs. caudal (increase) NAc-projecting vHPC neurons in mice making consummatory food port entries (29). By extension, it may be stipulated that in the present study, a reduction in vCA1 input led to an increase in avoidance bias via *caudal* shell mediated mechanisms, consistent with our previous work in which a pharmacological reduction in caudal NAc activity led to an avoidance bias in the face of motivation conflict. vCA1/subiculum inputs extend throughout the rostro-caudal axis of the NAc shell, and these projection neurons are not arranged topographically within the vHPC(29), hence future work is needed to address how differential afferent activation of the shell along the rostrocaudal axis is achieved to govern behavioral output.

In summary, these results demonstrate that the vCA1-NAc shell circuit is critically involved in biasing decision making towards approach under situations of motivational and social ambivalence, distinct from other circuits such as the vCA3 or vCA1-lateral septal circuits that promote avoidance behavior under conflict(16). Dysregulation of vCA1-NAc shell could underlie neuropsychiatric diseases by eliciting decision uncertainty which results in promoting approach in substance use disorders, while potentiating avoidance in mood disorders such as anxiety and depression (3,4,6,7,45).

## Materials and Methods

### Animals

63 adult male and female Long-Evans rats (Charles River Laboratories, Quebec) weighing 225-325g at the time of surgery were used in this study. Rats were pair-housed in a colony room maintained at a constant temperature (21°C) and humidity (35-60%) with a 12-hour light-dark cycle (light on at 7am). All behavioral testing, the order of which was counterbalanced between groups, occurred during the light hours and water was available *ad libitum*. Animals were food restricted for behavioral testing and were maintained at 85-90% of their pre-testing weight. All procedures were conducted in accordance with the regulations on animal experimentation established by the Canadian Council on Animal Care (CCAC) and approved by the Local Animal Care Committee at the University of Toronto.

### Surgery

Rats were anesthetized with isoflurane gas (Benson Medical, ON, Canada) and placed in a stereotaxic frame (Stoelting Co, IL USA). A midline incision was made, and the fascia retracted to reveal bregma. Rats were then micro-injected bilaterally with 0.5 μl of either the AAV8-CAMKII-hM4Di-mCherry or AAV8-CAMKII-EGFP control viruses (Addgene, MA) into the vCA1 (AP: -5.6, ML: ±5.8, DV: -7) over 5 min. After the viral injection, the injector remained in place for a further 10min for the AAV to diffuse away from the tip. Following AAV injection, bilateral cannulae (26 gauge; Plastics One, Roanoke, VA) were implanted into the NAc shell (AP: +1, ML: ±1, DV: -6.5) with the tips of the cannulae lying 1mm above the target site. The guide cannulae were secured to the skull with 4-5 jewellers’ screws embedded within dental cement. To maintain patency, stainless steel dummy cannulae were inserted into the guide cannulae. Rats were given, at minimum, 1-week post-surgery to recover, and the commencement of behavioral training was timed to ensure that the hM4Di receptors would have at least 6 weeks to express before activation.

### Drugs and Microinfusions

Clozapine-N-Oxide (CNO), an agonist for hM4Di, was mixed with physiological saline (0.9%) to achieve a concentration of 1mM. Prior to the first drug infusion, animals were acclimated to the infusion procedure. To achieve this, animals were sequentially exposed to the room, the restraint hold and the removal and insertion of the dummy cannula. On infusion test days, rats were bilaterally infused with 0.3 μL of either vehicle saline or CNO at an infusion rate of 0.3 μL/min via 30-gauge microinjectors connected to a 10 μl Hamilton syringe placed on an infusion pump (Harvard Apparatus, MA). Post-infusion, the infusers were left in place for an additional minute to allow for drug diffusion. Animals were then given a 10-minute adjustment period prior to behavioral testing.

### Mixed Valence Y Maze task

#### Apparatus

Behavioral training and testing occurred in a six-arm radial maze apparatus (Med Associates, USA). Access to the arms (45.7 cm (L), 9 cm (W), 16.5 cm (H)) was controlled by automated guillotine-like doors. Only three of the six arms (forming a Y-maze) were used at any one time. Each arm was enclosed with plexiglass walls, a removable lid and contained a steel grid floor. The walls and lid were covered with red cellophane to obscure extra-maze stimuli and the steel grids were connected to a foot shock generating device (Med Associates, USA). Each arm also contained a receding well at the end, which was connected to a syringe via polyethylene tubing to allow the delivery of a liquid sucrose solution (20%). The maze was wiped down with 70% ethanol solution after each training session to eliminate odor traces, and rotated by varying degrees (60^°^, 120^°^, or 180^°^) at the end of each day to minimize conditioning to intra-maze cues. A ceiling-mounted camera was used to record test sessions.

### Cues

During training and testing, three sets of visuotactile bar cues with different textures, colours and reflective properties (45 cm x 2.5 cm) were attached to the lower portion of the maze arms. Each bar insert became associated with either sucrose delivery (appetitive), mild foot shock administration (aversive) or no outcome (neutral). The valence of cues for each rat was determined following the unvalenced cue habituation session (described below). This allowed us to counterbalance any innate preferences for a cue by assigning the opposite valence during conditioning.

### Habituation

As described in detail previously (14,16,27,32) rats received 4 separate 5min habituation sessions. During the first session, rats were habituated to the Y-maze apparatus without the insertion of any bar cues. The rats were first placed in the central hub. After a minute, the doors leading to all three arms were raised and the rats were allowed to freely explore the maze for 5 minutes. During the second session, the visual-tactile cues were inserted into the arms. Rats then freely explored the maze and cues for 5 minutes. Their time spent in each arm, each with a different cue, was recorded and used to assign cue valence. Specifically, the least preferred cue was assigned as the appetitive cue, the most preferred cue as the aversive cue, and the remaining cue as the neutral cue. During the third session, rats were presented with two sets of cues in two arms. One arm contained a combination of the to-be-assigned appetitive and aversive cue while the other contained the neutral cue. This ensured that the combination of these two cues, which occurs during the final conflict test, was not a novel stimulus. During the final habituation session, rats were placed into the central hub for 30 seconds. They were then confined in each of the three un-cued arms, separately, for 2 minutes before being returned to the hub.

### Cue Conditioning

Animals underwent 9-12 conditioning sessions to acquire appetitive and aversive cue conditioning. Each session began with 30 seconds in the hub, followed by 2 minutes in each of the individual arms. In the arm containing the appetitive cue, rats received 4 x 0.4 mL, 20% sucrose delivered at random intervals averaging 30s. In the arm containing the aversive cue, rats received 4 mild foot shocks (1 s, 0.25-0.30 mA) delivered at random intervals. In the arm with the neutral cue, no outcomes were given. The order of arm presentation was varied daily to prevent animals associating the outcomes with the order of presentation. Additionally, arm cue placement was counterbalanced between animals and varied between sessions. The maze was also rotated daily to limit the effect of any extra-maze stimuli.

### Conditioned Cue Acquisition Test

After every fourth conditioning session, rats underwent a conditioned cue preference/avoidance test to assess their learning. Rats freely explored the cues in the maze for 5 minutes in the absence of sucrose or shock. Time spent in each arm was measured. Successful acquisition of conditioned cue preference and avoidance was indicated by rats spending more time in the appetitive arm than the neutral and aversive arms (conditioned cue preference) and spending the least amount of time in the aversive arm (conditioned cue avoidance).

### Mixed Valence Conflict Test

Before being placed into the radial maze for the mixed valence conflict test, rats received either a CNO or saline microinfusion, as described above. During this test, rats freely explored two arms for 5 minutes. One arm contained both the appetitive and aversive cues while the other arm contained only the neutral cue. We recorded multiple behavioral measures as indices of approach/avoidance, including the time spent in each cued arm and the hub, latency to enter each arm for the first time, the number of full body entries in to the arms, retreats (half body entries into the arm followed by rapid withdrawal into the hub, or back treading once inside the arm) and risk assessment behavior (number of times spent looking/checking back and forth between arms before making their first entry into an arm).

### Cue Preference and Avoidance Tests

Following the conflict test, animals underwent a refresher cue conditioning session. A day later, rats received a drug (CNO or saline) microinfusion as described above, and underwent cue preference and cue avoidance tests, the order of which were counterbalanced across animals (and conducted a minimum of 30min apart). During these tests, rats freely explored two of the arms for 5 minutes in extinction. One arm contained either the appetitive or the aversive cue while the other arm contained only the neutral cue. Total time spent in each arm, hub, as well as other indices of approach/avoidance behaviors were measured.

### Cue Novelty Test

A subgroup of animals (n=12) received another refresher cue conditioning session, and were then administered a cue novelty test prior to which they received a CNO microinfusion in a between-subject design as the novelty test cannot be repeated (n=6 hM4Di, n=6 GFP). In this test, animals were allowed to freely explore two arms of the Y-maze for 5 minutes, this time with one arm containing a novel set of cues, and another containing the neutral cues. Total time spent in each arm and hub, as well as other indices of approach/avoidance behaviors were measured.

### Social Preference and Memory Test

We used a social discrimination paradigm to test the effects of vCA1-NAc shell inactivation on social preference and memory. Rats were habituated to a circular open arena [136 cm (D), 75cm (H)] with two empty wire cages [25 cm (L) x 20 cm (W) x 15 cm (H)] placed 50cm apart for 15min. Following the habituation session, rats received a microinfusion of either saline or CNO, and were then placed into the centre of the arena to explore the apparatus and allowed to interact with a caged conspecific and an empty cage for 10 minutes before being removed. After a 20-minute delay period, rats were placed back into the area, but this time with two caged conspecifics to interact with (one novel and one familiar). In a small subset of animals (n=8), the rats were presented with two caged conspecifics (instead of one) in the first phase. However, this version was soon abandoned for the remaining animals as we wished to utilize the first phase as a sociability test (preference for social stimulus over no stimulus). Thus, only data from the remaining animals are reported. All tests were recorded using a video camera and the time spent interacting with and sniffing each cage was measured.

### Locomotor Test (LM)

After a saline or CNO microinfusion, all animals were placed into individual cages (44 cm [L] x 24 cm [W] x 20 cm [H]) with standard bedding material and stainless-steel cage lids. Animals were left for 1 hour while their locomotor behavior was monitored using an overhead camera and processed by ANY-maze tracking software. Total distance traveled (in m) was recorded in 5-minute time bins.

### Elevated Plus Maze (EPM) Test

An elevated plus maze task was used to measure unconditioned anxiety levels. The apparatus was placed in a novel room and was elevated 50 cm from the floor. It contained a central hub [10 cm (L) x 10 cm (W)] that connected four arms [40 cm (L) x 10 cm (W)], two of which were enclosed by walls [22 cm (H)]. After infusion of either saline or CNO, rats were placed in the central hub of the maze facing an open arm and given 10 minutes to explore the maze. Time spent in the open and closed arms as well as the middle area were measured. Additionally, the number of entries into each arm was recorded.

### Open Field Test

An open field test was conducted as a final test prior to euthanasia, to endogenously activate c-FOS labeling in the region of interest for immunohistochemical analysis. Rats were placed in an open field arena [45 cm (L) x 45cm (W) x 40 cm (H)] for 10 minutes. This apparatus consisted of four black opaque walls and the floors were lined with fresh bedding. The arena was divided into a 3 x 3 grid, and the time spent in the center zone (grid in the centre, 15 x 15 cm) and periphery of the apparatus was analyzed using NOLDUS EthoVision XT software.

### Histology

Rats underwent transcardial perfusions with 4% paraformaldehyde (PFA) within 75-90min after the completion of the open field test, and their brains were extracted. The brains were sliced into 50 μm coronal sections using a vibratome (Leica VT1200S). A subset of these slices was then analyzed to confirm viral expression and cannula placement.

### c-Fos Immunohistochemistry

Brain sections were treated with 1% hydrogen peroxide, then incubated in 0.5% 1,3,5-Trinitrobenzene (TNB) blocking solution for 1 hr at room temperature and incubated overnight at 4 °C with rabbit c-fos antibody (1:5000 dilution, Synaptic Systems, Goettingen, Germany). Following incubation with the primary antibody, sections were washed in PBS (3×5 min) and incubated overnight with secondary antibody (peroxidase conjugated donkey-anti-rabbit 1:500, Jackson Immunoresearch, Baltimore, PA, USA). The Tyramide Signal Amplification procedure was then employed, using NHS-fluorescein for the hM4Di-expressing brains, or NHS Rhodamine for the GFP control brains (Thermo Fisher Scientific, MA) as dye substrates. After a final round of PBS washes, brain slices were and mounted on gelatin-coated slides and air dried before being coverslipped with Fluoroshield Mounting medium with DAPI for nuclear staining (Abcam, MA).

### Cell imaging and counting

hM4Di and GFP expression, and c-Fos immunoreactivity were subsequently visualized at 4x, and 20x magnification using the NIKON Ti2-E microscope (NIKON, NY). GFP expressing cells and c-Fos positive cells conjugated with TSA-fluorescein were visualized using the FITC filter (excitation: 467 - 498nm; emission: 513 – 556 nm), while hM4Di expressing cells and c-Fos positive cells conjugated with TSA-Rhodamine were visualized using the TexRed filter (excitation: 532 – 587 nm; emission: 608 – 683 nm). Quantification of c-fos positive cells was achieved using two images of coronal sections of the NAc taken at 4x magnification from each animal. Projections from the ventral CA1 to the shell were demarcated as the regions of interest, and the number of cfos-positive cells within those boundaries were counted by converting the images into 8-bit and using ImageJ software (Rasband, W.S., U. S. National Institutes of Health).

### Statistical Analysis

All data collected were analyzed using SPSS (v25) and graphed in GraphPad Prism version 10.0.2 (GraphPad Software, La Jolla, CA, USA).

To assess the acquisition of cue associations during the training period, acquisition data (time spent in each arm) were subjected to a repeated measures analysis of variance (ANOVA) test with Arm (Appetitive, Neutral, Aversive) as a within-subjects factor. To determine whether there were any preexisting group differences, Sex (Male, Female), Virus Condition (GFP, hM4Di) and future Drug Condition (Saline, CNO) were used as between-subjects factors. Data from the Y-maze conflict test and cue preference/avoidance tests (time spent in each arm, latency to enter, ‘risk assessment’ behavior, number of entries and retreats) were subjected to a repeated measures ANOVA with Sex, Drug and Virus as between-subjects factors and Cue (Conflict, Neutral; Appetitive, Neutral; Aversive, Neutral) as a within-subjects factor. Data from the novelty cue preference were also subjected to ANOVA with Virus as a between-subject factor and Cue (Novel, Neutral) as a within-subject factor. For all tests, central hub time was analyzed separately using ANOVA with Virus, Drug and Sex as between-subject factors.

Social data (interaction time) were analyzed with Virus, Drug and Sex as between-subjects factors and Stimulus (Novel vs no conspecific for first social preference test) or Novelty (Familiar, Novel for social memory test) as a within-subjects factor.

Locomotor data (distance travelled) were split into twelve 5-minute time bins and analyzed using a repeated measures ANOVA with Sex, Virus and Drug as between-subjects factors and Bin (1–12) as a within-subjects factor. Similarly, both anxiety tests (elevated plus maze and open field) were subjected to repeated measures ANOVA. For EPM, time spent in each arm was analyzed with Virus, Drug and Sex as between-subjects factors and Arm (Open, Closed) as a within-subjects factor. For open field test data, time spent in the central grid was analyzed by independent samples t-tests.

Finally, the density of cfos positive cells in the NAc shell of the hM4Di virus group was analyzed using a 2-way ANOVA with Drug (Saline, CNO) and Sex as between subject factors.

All tests were accompanied by tests of homogeneity of variance and sphericity, and significant main/interaction effects were followed up with planned comparisons and post-hoc tests where appropriate, using Bonferroni correction to control for Type 1 error in multiple comparisons. The alpha level was set at p < 0.05, and Bonferroni corrected p-values are reported.

## Acknowledgements

This study was funded by a project grant from the Canadian Institutes of Health Research awarded to RI & ACHL (156070) and a Canada Graduate Scholarship awarded to DP by Social Sciences and Humanities Research Council (SSHRC).

